# The tRNA pseudouridine synthase TruB1 regulates the maturation and function of let-7 miRNA

**DOI:** 10.1101/2020.02.16.951954

**Authors:** Ryota Kurimoto, Tomoki Chiba, Yoshiaki Ito, Takahide Matsushima, Yuki Yano, Kohei Miyata, Tsutomu Suzuki, Kozo Tomita, Hiroshi Asahara

## Abstract

Let-7 is an evolutionary conserved microRNA that mediates post-transcriptional gene silencing to regulate a wide range of biological processes, including development, differentiation, and tumor suppression. Let-7 biogenesis is tightly regulated by several RNA-binding proteins, including Lin28A/B, which represses let-7 maturation. To identify new regulators of let-7, we devised a cell-based functional screen of RNA-binding proteins using a let-7 sensor luciferase reporter, and identified the tRNA pseudouridine synthase, TruB1. TruB1 enhanced maturation specifically of let-7 family members. Rather than inducing pseudouridylation of the miRNAs, HITS-CLIP (High throughput sequencing crosslinking immunoprecipitation) and biochemical analyses revealed direct binding between endogenous TruB1 and the stem-loop structure of pri-let-7, which also binds Lin28A/B. TruB1 selectively enhanced the interaction between pri-let-7 and the microprocessor DGCR-8, which mediates miRNA maturation. Finally, TruB1 suppressed cell proliferation, which was mediated in part by let-7. Altogether, we reveal an unexpected function for TruB1 in promoting let-7 maturation and function.

## Introduction

MicroRNAs (miRNA) are short non-coding RNAs of about 22 bases that regulate gene expression by inhibiting either the stability of target mRNAs or protein synthesis in association with Argonaute (AGO) family proteins (Lee et al., 1993; Wightman et al., 1993). Lin-4 and let-7 were the first reported miRNAs in *C. elegans* (Lee et al., 1993). Since then, more than 1,500 conserved miRNAs have been identified across many species including mammals (Ambros., 2004). The expression and maturation of miRNAs are tightly regulated by multi-step mechanisms involving protein-RNA associations with enzymatic activities, and molecular shuttling, specifically: i) Pol-II mediated primary miRNA (pri-miRNA) transcription (Lee et al., 2004; Cai et al., 2004), ii) Drosha/ DiGeorge Syndrome Critical Region 8 (DGCR8)-mediated cleavage processing producing a precursor-miRNA (pre-miRNA) in the nucleus (Denli et al., 2004; Gregory et al., 2004; Han et al., 2004; Landthaler et al., 2004; Lee et al., 2003; Zeng et al., 2005), iii) nuclear export of pre-miRNA by Exportin-5 (Yi et al., 2003; Bohnsack et al., 2004; Lund et al., 2004), iv) additional cleavage by the RNase III enzyme Dicer in the cytoplasm (Bernstein et al., 2001; Grishok et al., 2001; Hutvágner et al., 2001; Ketting et al., 2001; Knight and Bass, 2001), and v) incorporation into the RNA-induced silencing complex (RISC) with AGO family proteins to generate the final mature miRNA (Hammond et al., 2001; Mourelatos et al., 2002; Tabara et al., 1999). Although the first transcription step for pri-miRNA expression can be driven by transcription factors and RNA Pol-II as well as mRNA, the following maturation steps, ii-v), are also critical and unique to miRNA biogenesis and dynamics.

The comparison of gene expression in tumor tissues showed discrepancies in the levels of mature miRNAs and their pri-miRNAs (Thomson et al., 2006). In particular, mutually exclusive and reciprocal regulation between the let-7 miRNA and the RNA binding protein (RBP) Lin-28 at these maturation process have been well demonstrated in ES cells, *C.elegans*, and cancer pathology (Newman et al., 2008; Piskounova et al., 2008; Heo et al., 2008; Viswanathan et al., 2008; Chang et al., 2009). Lin28A/B specifically binds to the preE loop sequence of pri-/ pre-let-7 and suppresses the processing of let-7 by causing oligo-uridylation of the let-7 precursor through TUT4/7 activity (Heo et al., 2008; Piskounova et al., 2011). Based on these fundamental findings, the factors mediating let-7 multi-step regulation, which may be critical for various biological and pathological events, have been extensively explored by several approaches (Newman et al., 2008; Trabucchi et al., Treiber et al., 2017); however, to date, only a few factors have been shown to regulate let-7 family-specific maturation (Newman et al., 2008; Trabucchi et al., Michlewski et al., 2010; Choudhury et al., 2014; Pandolfini et al., 2019).

Taking advantage of the completion of genome-wide comprehensive full-length cDNA libraries (Iourgenko et al., 2003; Chanda et al., 2003; Conkright et al., 2003; Huang et al., 2004; Liu et al., 2005), we and others successfully developed cell-based screening systems to quantify miRNA targets and functions, including destabilization of target mRNA and suppression of protein synthesis (Wolter et al., 2014: Wolter et al., 2015; Ito et al., 2017). Here we applied the above strategy and introduced a new cell-based screening to identify genes that regulate let-7 miRNA by using a luciferase reporter assay with a set of expression plasmid libraries mainly encoding proteins with RNA binding properties. We identified a new regulatory mechanism: TruB1, an RNA-modifying enzyme, selectively regulates let-7 activity in an enzymatic activity-independent manner.

## Results

### Functional screening to detect RNA binding proteins (RBPs) that promote let-7 expression

We developed a cell-based, functional screen using an expression plasmid library and a let-7 sensor reporter to identify proteins that regulate let-7 biogenesis. This type of screen enables the identification of proteins that regulate endogenous let-7 function, as opposed to other aspects of let-7 biology, such as binding, which is assayed using affinity-based (pull-down) screens. The gain-of-function format also enables the identification of regulatory factors that may cause lethality under loss of function conditions, such as with Crispr-Cas9 or RNAi-KD screens. The let-7 sensor reporter was designed to monitor let-7 expression levels by inserting a let-7a target site into the luciferase 3’UTR region driven by the SV-40 promoter in pLuc2 plasmids (Fig 1 A). Thus, luciferase activity is repressed in the presence of mature let7 miRNA (Fig EV1). In this screen, we focused on identifying molecules directly regulating the miRNA maturation process, thus we prepared a sib-selection library covering molecules annotated with RNA binding protein (RBP) features from the Center for Cancer systems Biology (CCSB)-Broad Lentiviral over-expression library (Yang et al., 2011). As many zinc-finger-proteins have not been directly tested for their RNA or DNA binding preference, the genes with zinc-finger domains were also included in the sib-selection library. Ultimately, 1469 genes were selected and prepared for the screening. This library was transfected into HEK293FT cells, along with the let-7 sensor reporter. If let-7 maturation is inhibited by coexpression of one of the cDNAs in the RBP library, expression of the luciferase reporter would be induced. As a positive control for the assay, we over-expressed Lin28A from the same backbone plasmid transfection along with the let-7 sensor reporter with or without overexpression of pri-let-7a in HEK293FT cells. This significantly promoted luciferase activity reflecting reduction of endogenous mature let-7 expression (Fig. 1B). Next, we screened the 1,469 genes in our library utilizing the luciferase assay with the let-7 sensor reporter in HEK293FT cells seeded into 384 well plates. pRL-SV40 Renilla luciferase activity was used as a transfection efficiency control. We only included genes that did not change the luciferase activity of the let-7 sequence minus the reporter (0.5 <, > 2.0) (results are shown in Appendix Table S1). Of these, we found that over-expression of 259 genes significantly reduced relative luciferase activity (< 0.50)compared with the GFP expression plasmid control (Fig. 1D), indicating that they potentially promote let7 miRNA maturation. From the top five hits, we performed a larger scale (96 well plate) luciferase reporter assay for validation, whereby four of the top five genes reproducibly repressed the luciferase activity of the let-7 sensor (Fig. 1E). To confirm the effect on endogenous let-7a expression, we examined the expression levels of a panel of ten miRNAs including let-7a by qPCR. Consistent with the screening results, overexpression of all of the top five candidate genes significantly promoted endogenous mature let-7a expression. Only TruB1 selectively induced let-7 expression but did not increase expression of the other miRNAs, whereas the other four hits, SF3A3, LARP7, GLTSCR2, and EF1E1, also increased the other miRNAs (Fig 1F). This suggests the specific promotion of let-7 biogenesis by TruB1. TruB1 is among the RNA-modifying enzymes that mediate pseudouridylation of tRNA. TruB1 has also recently been shown to regulate mRNA via pseudouridylation (Schwartz et al., 2014; Safra et al., 2017), but, to date, there has been no reported function of TruB1 in miRNA biogenesis.

**Figure 1.**
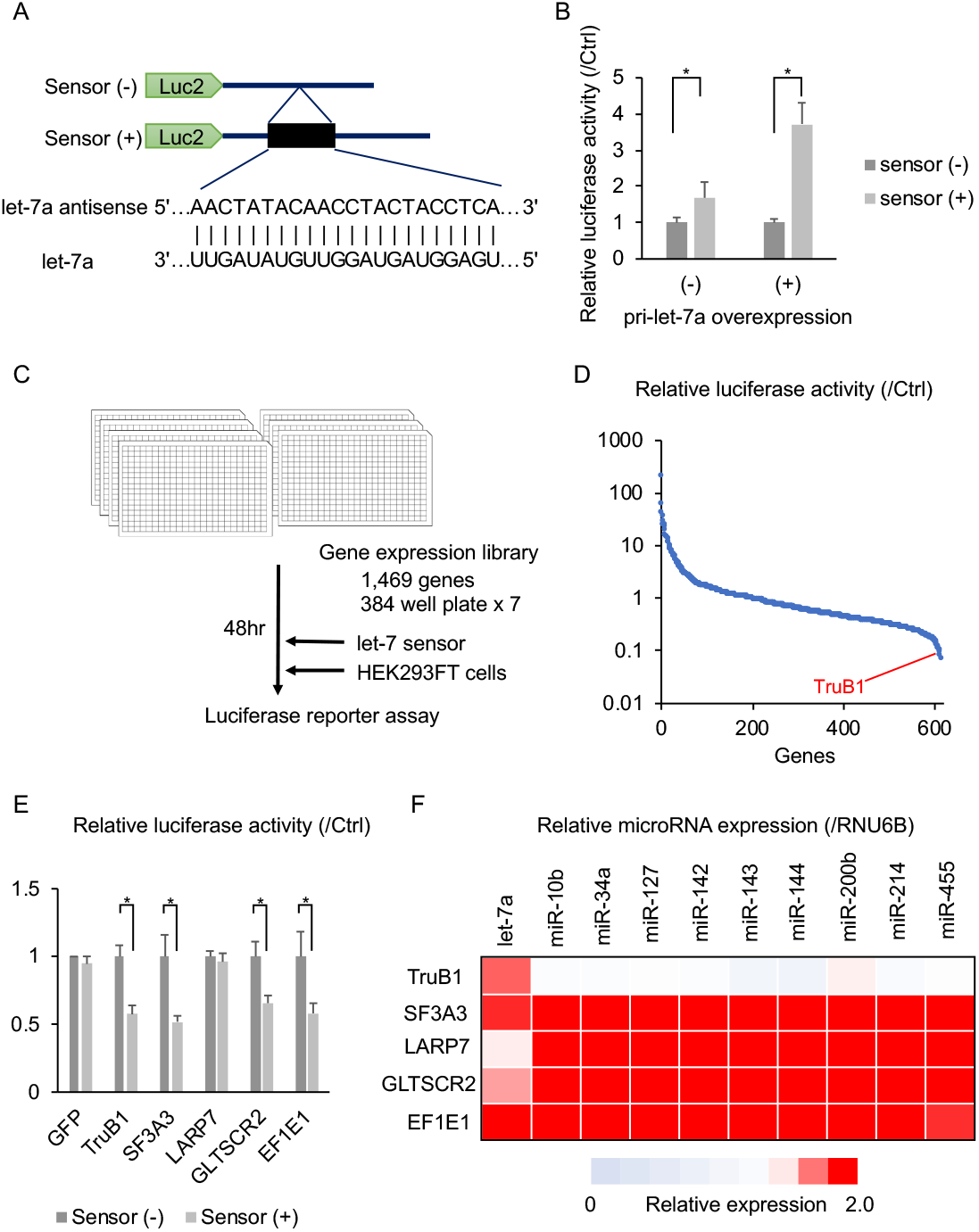
Functional screen using a luciferase reporter assay to identify RBPs inducing let-7-expression. (A) Design of the let-7 sensor reporter and sequence of let-7a. Sensor (+) vector has the antisense sequence of let-7a inserted into the 3’UTR of the luciferase gene. (B) Relative luciferase activity of the let-7 sensor reporter (sensor (+)) or the negative control sensor (−) reporter contransfected with pLX-Lin28A or the GFP vector (Ctrl) in HEK293FT cells with or without pcDNA-pri-let-7a transfection. Error bars show SD; n=3. (C). The screening model. A screening in 384-well plates with a library of 1469 genes was performed. (D). Screening results: relative luciferase activity of each gene in the library. Only genes that did not affect the luciferase activity of the sensor (−) reporter (0.5 <, > 2.0) were analyzed. The full results of the screen are also in Appendix TableS1. (E). Relative luciferase activity of the sensor (+) reporter or sensor (−) reporter induced by expression vectors of the top 5 genes identified in the screen or the GFP vector in HEK293-FT cells. Error bars show SD; n=3. (F). Heat map of relative miRNA expression of various miRNAs in HEK-293FT cells transfected with expression vectors of the top 5 genes from the screen, or the ctrl (GFP) vector. Red color represents suppressed expression compared with ctrl (GFP). n=3.

### TruB1 selectively promotes the expression of let-7 families

Although a few proteins, such as Lin28A/B, KSRP, hnRNPA1, have been shown to be involved in the biogenesis of a specific set of miRNAs, most other miRNA regulators act without specificity, or their miRNA selectivity remains unclear. To test whether TruB1-dependent let-7 promotion is specific for the let-7 family, as our preliminary results suggested, we comprehensively analyzed the effect of TruB1 knockdown by siRNA on miRNA biogenesis in HEK293FT cells by TaqMan array. TruB1 knockdown down regulated the expression of five mature let-7 family members (let-7a, b, c, f and g) (Fig 2A, Fig EV2A, Appendix Table S2). These data were further confirmed by large scale TaqMan PCR for miRNA and northern botting (NB), revealing that all nine endogenous mature let-7 family genes were significantly downregulated upon TruB1 knockdown(Fig 2B-D). In contrast, TruB1 knockdown increased the expression levels of endogenous, immature, primary-let7 (Fig 2E). We confirmed these findings in both A549 and HeLa cells (Fig EV2B,C). These results indicate that TruB1 promotes let-7 family miRNA biogenesis specifically at their maturation step, rather than at the transcriptional level.

**Figure 2.**
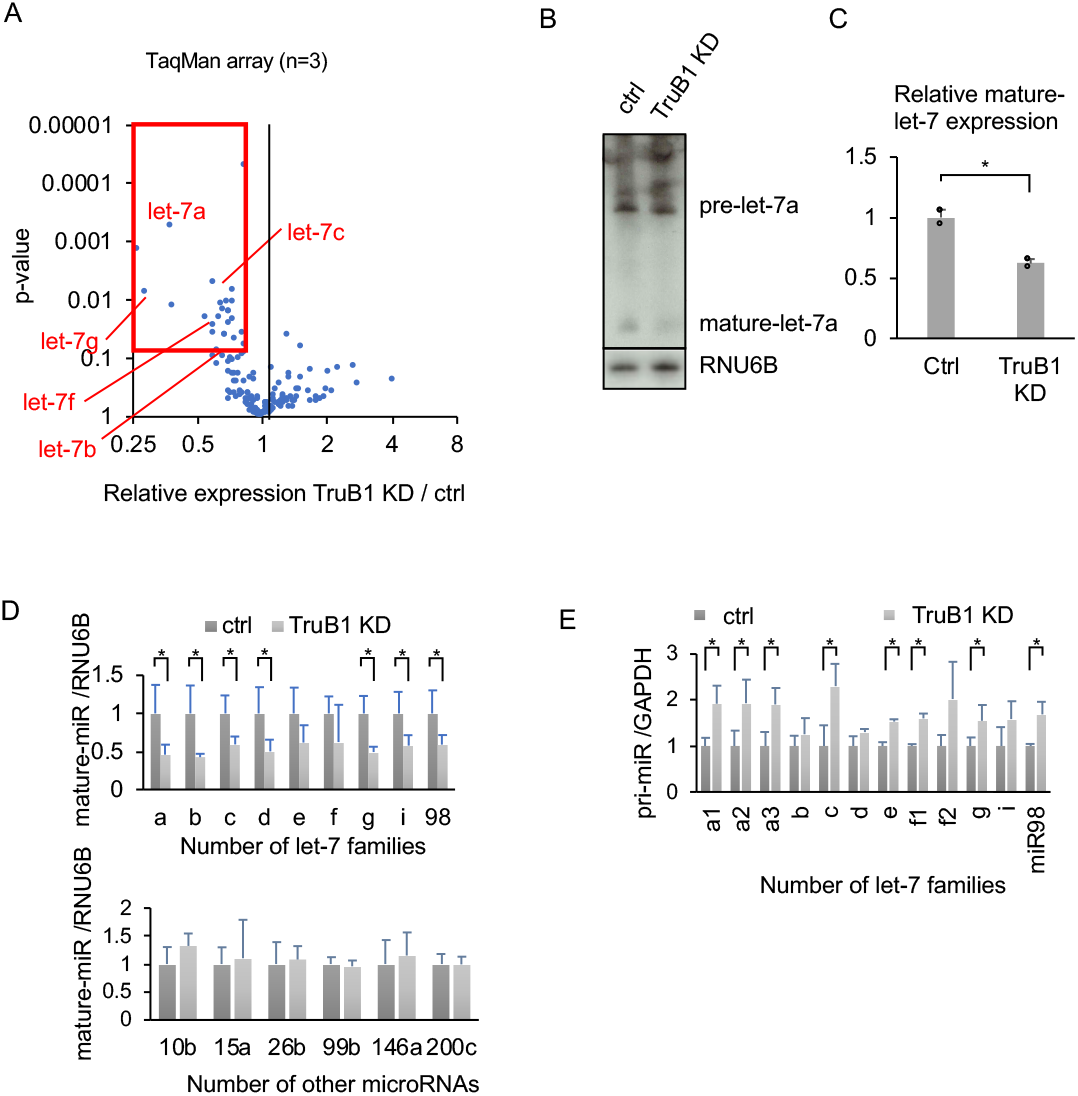
TruB1 selectively promotes the expression of let-7 family miRNAs. (A). Volcano plot of TaqMan array showing the mean average expression and p-value of miRNA expression profiles between TruB1 KD and ctrl (scramble). N=3. MiRNAs significantly suppressed are enclosed by the red square. (B). Northern blotting for let-7a and RNU6B in HEK293FT cells with TruB1 KD or ctrl (siRNA). (C). Hybridization intensities of (B) were quantified and normalized to ctrl. (D). Relative expression of let-7 family miRNAs and other miRNAs in HEK293FT cells with TruB1 KD or ctrl (siRNA) determined by qRT-PCR. (E). Relative expression of primary-let-7 family miRNAs as in (D) determined by qRT-PCR. All experiments were performed in triplicate. Error bars show SD; n=3.

### TruB1 promotes miRNA processing independently of its enzymatic activity

Therefore, we analyzed whether let-7 regulation by TruB1 is dependent on the pseudouridine enzyme activity of TruB1. Critical residues involved in enzymatic activity (D48, D90) and RNA binding ability (K64) have been identified in *E. coli* TruB (Wright et al., 2011; Friedt et al., 2014; Keffer-Wilkes et al., 2016). Given the TruB amino acid sequences for these enzyme activity and RNA binding ability are highly conserved (Zucchini et al., 2003), we were able to insert mutations to modify the function of TruB1 by amino acid substitution. We generated two TruB1 mutants: Mt1, with inactivated enzyme activity, and mt2, with suppressed RNA binding ability (Fig 3A). An *in vitro* enzymatic activity assay with a tRNA^phe^ substrate using the wt, mt1 and mt2 recombinant proteins showed that the pseudouridylation enzyme activities of both mt1 and mt2 were completely attenuated (Fig 3B). An EMSA study using recombinant proteins revealed a physical interaction between tRNA^phe^ and wt TruB1 and mt1, but not with mt2 (Fig EV3A). Next, we tested the function of these mutants in cells. Western blotting (WB) revealed that all proteins were overexpressed largely to the same levels (Fig EV3B). Overexpression of Wt and mt1 TruB1 significantly increased let-7a maturation, whereas mt2 did not (qPCR, NB). These results suggest that promotion of let-7 maturation by TruB1 is independent of its enzyme activity, but dependent on RNA binding (Fig 3C-E).

**Figure 3.**
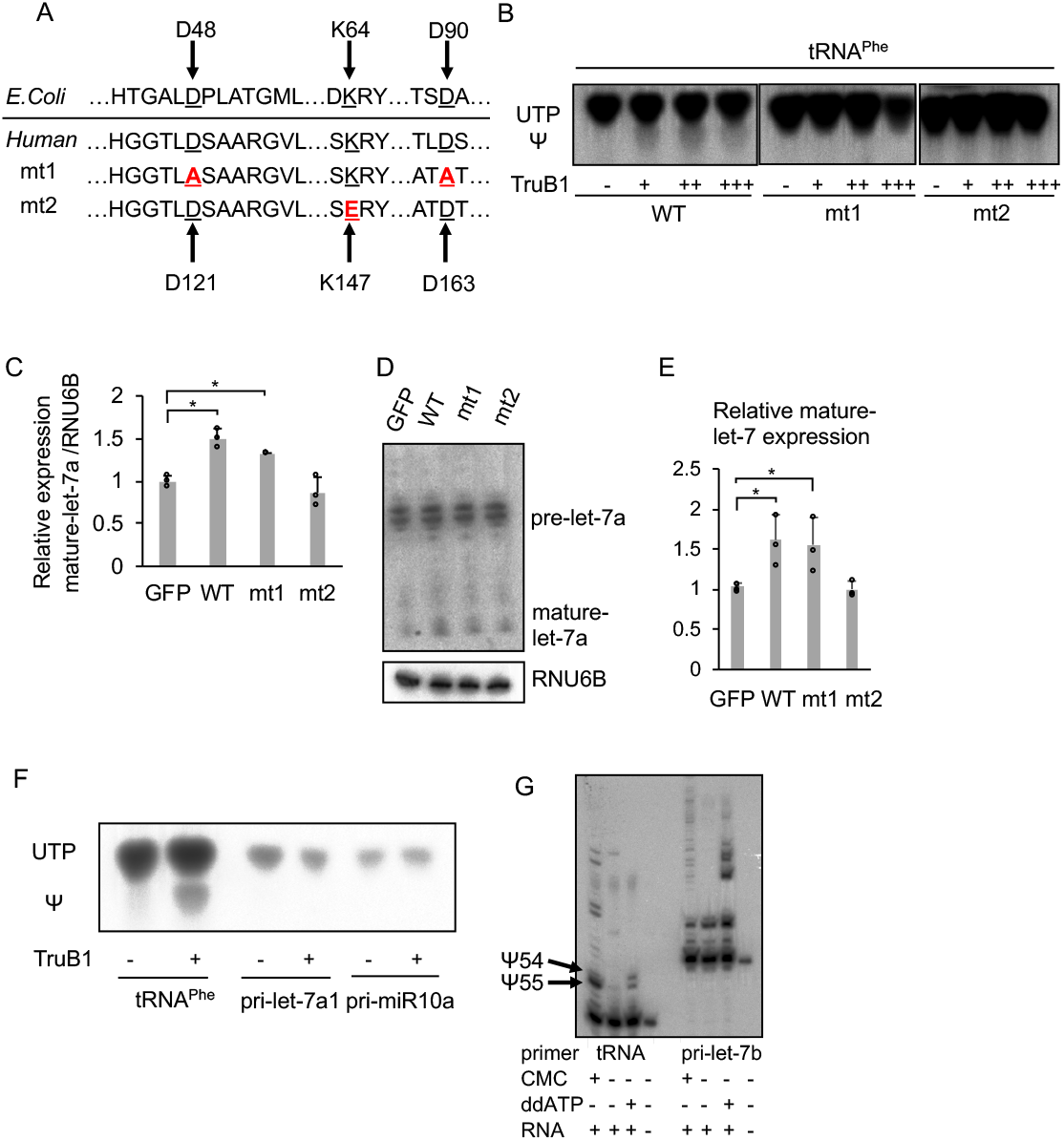
TruB1 promotes let-7 processing independently of its enzymatic activity. (A). Amino acid sequences encoding for the enzyme activity and RNA binding ability in *E.coli* TruB and human TruB1 (Top). Design of mutant 1 (mt1) and mutant 2 (mt2) (bottom). (B). In vitro enzyme activity assay. ^32^p-UTP-labelled tRNA^phe^ were treated with recombinant TruB1, mt1, or mt2. The strong upper bands represent UTP, and the lower weaker bands represent pseudouridine (Ψ) on autoradiographs of the TLC plate. (C). Relative miRNA expression of let-7a in HEK-293 cells infected with tetracycline-inducible expressing lentiviruses for TruB1, mt1, mt2 or GFP 5 days after doxycycline treatment, as determined by qRT-PCR. (D). Northern blotting for let-7a and RNU6B in HEK-293 cells infected with lentiviruses encoding tetracycline-inducible expression of TruB1, mt1, mt2, or GFP, 5 days after doxycycline treatment. (E). Hybridization intensities of (D) were quantified and normalized to ctrl (GFP). (F). Pseudouridylation activity of TruB1 for tRNA and pri-miRNAs. ^32^p-UTP-labelled tRNA^phe^, pri-let-7a1 or pri-miR10a were treated with recombinant TruB1. Upper bands represent UTP, lower bands represent pseudouridine (Ψ) in autoradiographs of the TLC plate. (G). Location of pseudouridine sites detected by the CMC primer extension method. Total RNA purified from HEK-293FT cells were treated with CMC. CMC treated RNA were reverse-transcribed with RI-labelled specific primers for tRNA^phe^ or pri-let-7b. ddATP was used for sequence control. Pseudouridines are indicated by black arrows. All experiments were performed in triplicate. Error bars show SD; n=3.

Next, we performed an in vitro enzyme assay to verify that let-7 could not be pseudouridylated by recombinant TruB1. Pseudouridine of tRNA (Phe) was clearly increased in the assay, whereas pseudouridylation of pri-let-7 and another primary miRNA was not observed (Fig 3F). We also examined the presence of pseudouridine directly by treatment with CMC (N-cyclohexyl-N0 -(2-morpholi-noethyl)carbodiimide metho-p-toluenesulfonate) followed by a primer extension assay. Although we could detect known pseudouridine sites in the tRNA, no pseudouridine site was found in let-7 (Fig 3G, Fig EV3C). We also synthesized RI-labeled pri-let-7a1 in which pseudouridine was randomly introduced using UTP : pseudouridine at a ratio of 1:1. An in vitro processing assay using this labelled pri-let-7a1 showed that the presence of pseudouridine in pri-let-7a1 did not affect its processing (Fig EV3D,E). These results indicate that the regulation of let-7 by TruB1 does not depend on its enzyme activity.

### TruB1 binds directly to primary-let-7

To determine how TruB1 regulates let-7 processing, we first tested whether TruB1 binds to pri-let-7. RNA immunoprecipitation (RIP) assays revealed that overexpression of wt and mt1 TruB1 immunoprecipitated pri-let-7a1, whereas mt2 did not (Fig 4A). An EMSA study using recombinant proteins revealed a physical interaction between pri-let7a1 and wt and mt1 TruB1, but not with mt2 (Fig 4B). Furthermore, EMSA was performed using mutant RNA in which the loop structure of pri-let-7a1 was modified (loop mt). As a result, no binding to TruB1 was observed in the loop mt (Fig 4B, Fig EV4A). To comprehensively survey the RNA-protein binding property under physiological conditions, we performed high-throughput cross-linking immunoprecipitation (HITS-CLIP) using a cell line in which the 3xFLAG tag was knocked into the TruB1 N-terminal end by using the CRISPR/Cas9 system (Fig 4C, Fig EV4B-F). This revealed the expected, but not previously shown, direct interaction between TruB1 and its known substrate tRNA (positive control). We also observed a physical interaction between TruB1 and pri-let-7a1 (Fig 4D, Appendix Table S3). Direct binding sites were also found in other miRNA sequences, including miR-29b, miR-139 and miR-107 whose expression levels of their mature forms were also specifically decreased upon TruB1 KD in the TaqMan array (Fig 4E, Appendix Table S3). Looking at the entire mapped reads, mRNA, lncRNA, and miRNA were detected in addition to tRNA (Fig 4F). It has been reported that mRNA is modified by TruB1, which is consistent with this result (Schwartz et al., 2014; Safra et al., 2017). When a read mapped to tRNA was extracted and motif analysis was performed, the sequence was similar to the pseudouridylation site reported for *Saccharomyces cerevisiae* PUS4 (Fig 4G), which is homologous with TruB1 (Fig 4H) (Becker et al., 1997). Although reads mapped to let-7a1 bound mainly near the terminal loop, no sequence similar to the tRNA motif was found in the let7 loop sequence. tRNA and let-7 are similar in that they form a loop structure (Fig 4I). Thus, we considered that TruB1 binds to let-7 in a structure-dependent manner. Interestingly, the TruB1 binding site on pri-let-7a1 also contained the sequence (GGAG), which is a binding motif of the CCHC-zinc domain of Lin28A / B. This suggests that TruB1 and Lin28 both bind to the same stem loop structure on *let-7*, and may compete for binding.

**Figure 4.**
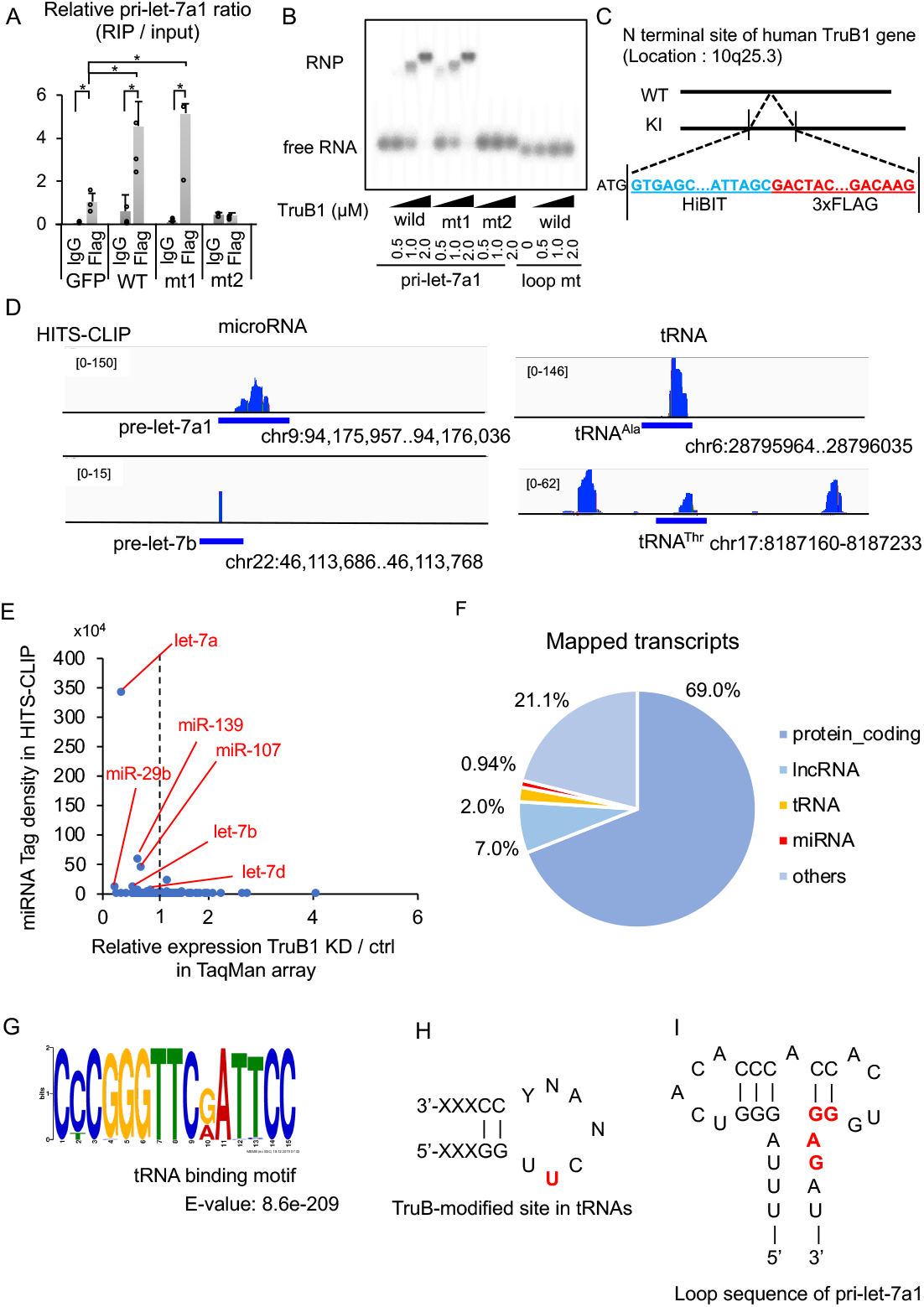
TruB1 binds to tRNA and primary-let-7. (A). RIP analysis of pri-let-7a1 and Flag-TruB1 from HEK-293FT cells. RNA was extracted from IP material and analyzed by qRT-PCR. HEK-293FT cells were infected with lentiviruses encoding tetracycline-inducible expression of TruB1, mt1, mt2, or GFP and treated with doxycycline. Error bars show SD; n=3. (B). EMSA of ^32^p-ATP-labelled pri-let-7a1 or pri-let-7a1 loop mt mixed with recombinant TruB1, mt1 or mt2 at several doses. RNP: Ribonucleoprotein complexes. (C). Design of Flag-labelled TruB1 knock-in (KI) cells. HiBIT and 3 x Flag sequences were inserted into the N-terminus of the TruB1 gene in HEK-293FT cells. (D) Sequencing clusters obtained from HITS-CLIP experiment. Left are for let-7, and right for tRNAs. (E). Tag densities represented as dots. Density of miRNA clusters in HITS-CLIP were normalized by miRNA expression as determined by TaqMan array. (F). The proportions of different types of mapped transcripts from the HITS-CLIP experiment. (G). Sequence motifs from reads mapped to tRNAs from the HITS-CLIP experiment. (H). TruB-modified site in tRNAs. Location of pseudouridine is colored in red. (I). Sequence of the terminal loop in pri-let-7a1. Lin28B binding site is colored in red.

### TruB1 enhances the affinity between microprocesser and primary let-7 and promotes microprocessing

The above results indicated that TruB1 directly binds to pri-let-7a1 under physiological conditions. To elucidate the function of this molecular interaction, we analyzed whether TruB1 was involved in the microprocessing step of miRNA biogenesis using an in vitro processing assay. RI-labeled pri-let-7a1 was incubated with a cell extract obtained from HEK293FT cells overexpressing different forms of TruB1. We found that the extract from cells overexpressing TruB1 promoted maturation of pri-let-7a1 into pre-/ mature-let-7a, whereas the extract from cells overexpressing mutant TruB1(mt2) did not. We also found that knockdown of endogenous TruB1 by siRNA inhibited maturation of pri-let-7a (Fig. 5A, B). These results suggest that let-7a biogenesis is promoted at the level of conversion from the pri-miRNA to the pre-miRNA. DGCR-8 is the microprocessor involved in the cleavage of pri-miRNA in the nucleus to form pre-miRNA. Thus, we tested the effect of TruB1 on the interaction between pri-let-7 and DGCR-8 by RIP affinity assay using an antibody to endogenous DGCR-8. Pri-let-7a1 was efficiently precipitated with DGCR-8, as expected. However, TruB1 KD by siRNA treatment reduced this interaction between pri-let-7a1 and DGCR-8 (Fig 5C), without affecting the expression level of DGCR-8. These data suggest that TruB1 is involved in enhancing complex formation between DGCR-8 and pri-let-7a1. Next, we tested the potential interaction between TruB1/let-7 and Lin28B, which also binds to the pri-let-7a1 stem loop region recognized by TruB1. In Lin28B knockdown cells, the ratio of TruB1-bound pri-let-7a1 over total pri-let-7a1 was increased, without altering TruB1 protein levels, as assessed by RIP (Fig 5D, Fig EV5). Thus, Lin28B inhibits the binding between TruB1 and pri-let-7a1.

**Figure 5.**
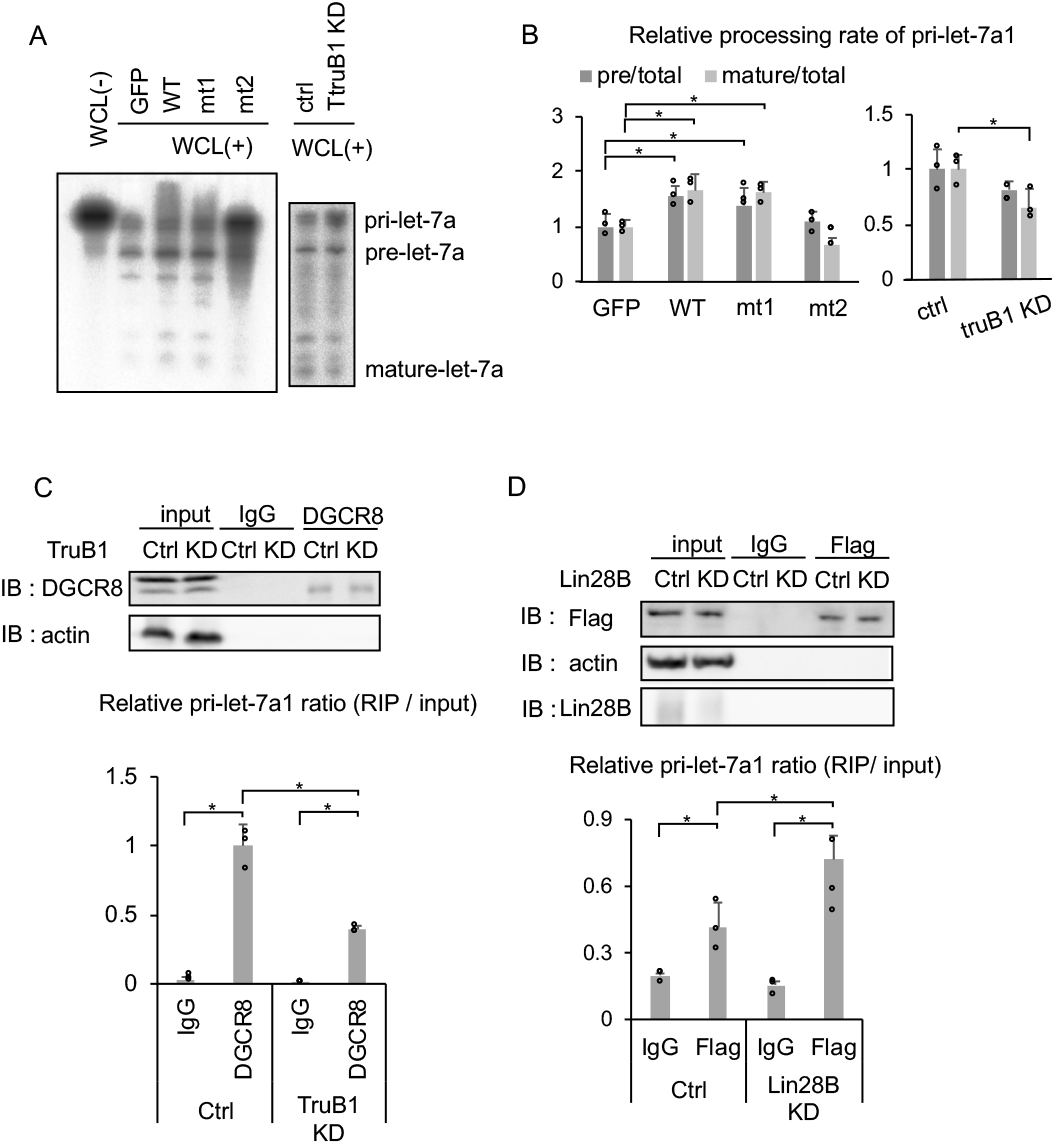
TruB1 promotes the microprocessing of primary let-7 and enhances binding of the microprocesser to primary let-7. (A). In vitro processing assay for RI-labelled pri-let-7a1. Autoradiographs of gels showing pri-let-7a1 treated with whole cell lysate (WCL) from HEK-293FT cells transfected with GFP, TruB1, mt1, or mt2 (overexpression, left), and TruB1 KD or ctrl (siRNA, right). (A). RI intensities of (B) were quantified and normalized to ctrl. Relative processing rate of pri-let-7a1 into pre- and mature- are shown. (C). RIP assay of pri-let-7a1 and DGCR-8 from HEK-293FT cells. Western blotting for input or immunoprecipitate (IP) using anti-DGCR-8 antibody and anti-actin antibody are shown on top. RNA was extracted from IP material and analyzed by qRT-PCR (bottom). (D). RIP assay of pri-let-7a1 and TruB1 from HEK-293FT cells with Lin28B KD or ctrl (siRNA). Western blotting for input or IP material using anti-Flag antibody, anti-Lin28B antibody, and anti-actin antibody are shown on top. RNA was extracted from IP material and analyzed by qRT-PCR (bottom). All experiments were performed in triplicate. Error bars show SD; n=3.

### TruB1 suppresses cell growth through the promoting let-7

Finally, we examined the cellular function of this new TruB1/let-7 axis by analyzing cell growth. Let-7 coordinates cell proliferation by targeting a set of genes, including KRAS, which is a proto-oncogene (Johnson et al., 2005; Wang et al., 2013). To test whether TruB1 can affect KRAS expression via let-7 regulation, we generated a Luc-reporter plasmid comprising the let-7 target region of KRAS in its 3’UTR, and performed a luciferase reporter assay (Fig 6A). As a result, overexpression of wt TruB1 suppressed the luciferase activity of the KRAS reporter, whereas mt2 did not (Fig 6B). Next, the effect of TruB1 overexpression on HEK-293FT cell growth was evaluated by real-time glo assay. Cell proliferation was significantly suppressed by TruB1 overexpression. To test whether this was mediated via let-7, we introduced RNAi to target the common seed sequence in the let-7 family. KD of the let-7 family partially rescued the suppressive effect of TruB1 overexpression on cell proliferation (Fig 6C, D, Fig EV6A). Taken together, these results indicate that cell proliferation can be regulated by the TruB1 and let-7 molecular cascade.

**Figure 6.**
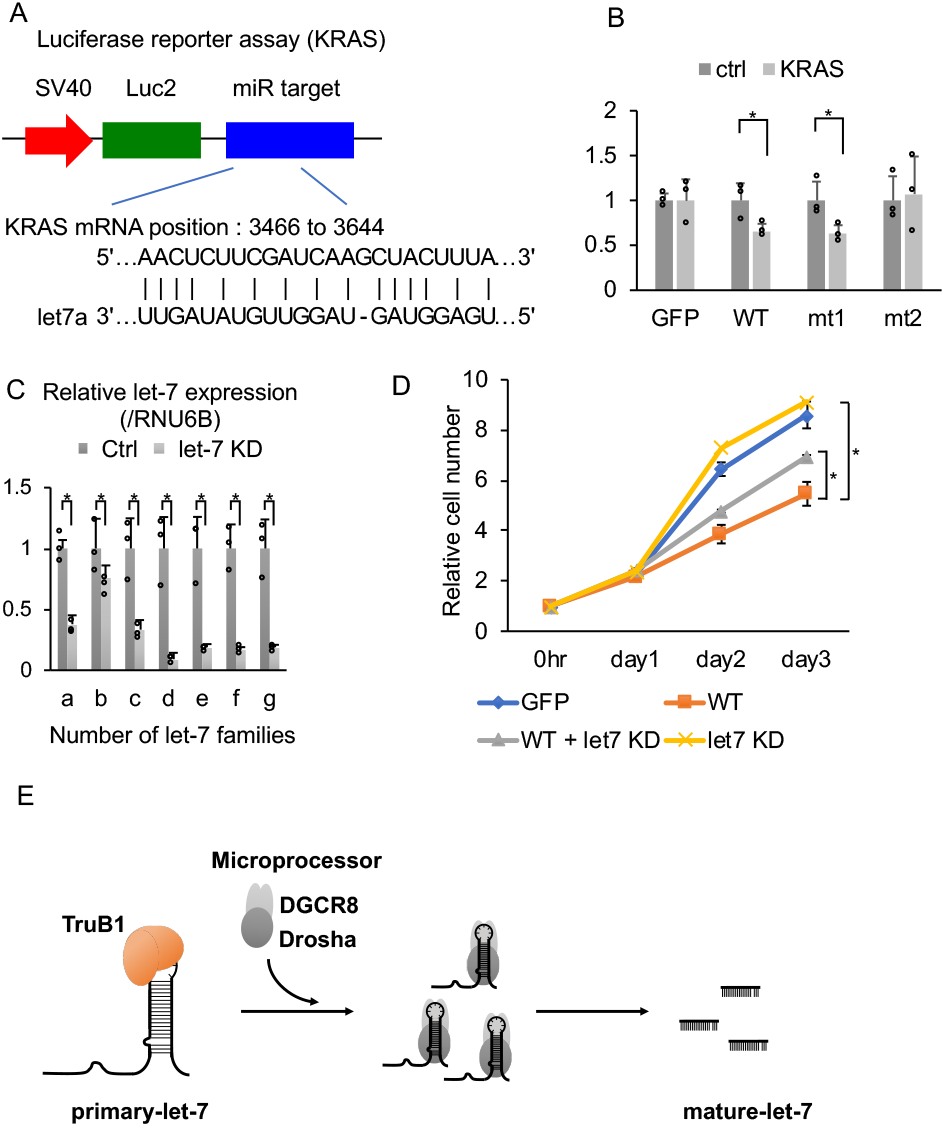
TruB1 suppresses cell growth by regulating let-7. (A). Design of luciferase reporter vector comprising the let-7 target region of KRAS mRNA (KRAS reporter). (B). Relative luciferase activity of KRAS reporter or Ctrl (empty) reporter in HEK293FT cells infected with lentiviruses expressing tetracycline-inducible TruB1, mt1, mt2, or GFP 5 days after doxycycline treatment. (C). Relative expression of let-7 families determined by qRT-PCR in HEK-293FT cells with KD of let-7 family members or ctrl (using 2’-O-methylated antisense inhibitor). (D). Real-time glo assay for HEK293FT cells infected with lentiviruses expressing tetracycline-inducible TruB1 or GFP, 5 days after doxycycline treatment, with or without KD of let-7 family members. (E). Schematic model for TruB1-dependent induction of let-7. All experiments were performed in triplicate. Error bars show SD; n=3.

## Discussion

In this study, we performed a systematic cell-based gain of function screen by combining an arrayed overexpression plasmid library and reporter system detecting mature let-7 expression and discovered a new function for TruB1, which is a pseudouridine RNA-modifying enzyme.

To date, various genes regulating miRNA maturation have been identified as co-immunoprecipitated factors from biotin-labeled miRNA hairpin, pri-miRNAs or microprocessors (Chendrimada et al., 2005; Lee et al., 2006; Auyeung et al., 2013; Treiber et al., 2017); however, only a few of them have shown selectivity for miRNA biogenesis. In this study, we could identify a set of molecules regulating let-7 biogenesis. Our strategy to use a cell-based and functional assay with an arrayed plasmid library has the following advantages: 1. The identified candidates are screened by their ability to regulate endogenous let-7 expression. Thus, the molecular function of the candidate proteins identified is already ensured, compared to a molecular affinity-based screening strategy, which only identifies binding. 2. The use of a prepared set of genes from the library enables us to perform a quantitative and more efficient screen.

All the raw data obtained in this approach is informative. 3. The gain of function approach can identify factors that are necessary for cell survival, whereas loss of function screens, such as those using shRNA or CRISPR/Cas9, may be unable to detect them. This is particularly relevant when performing screens with molecules performing fundamental functions, such as let-7.

In our approach, we identified a set of proteins that significantly enhance endogenous let-7 expression. Among them, TruB1 is shown to play a critical role in let-7 specific miRNA maturation, whereas the other candidates did not show such selectivity.

TruB1 is a pseudouridine synthase (Zucchini et al., 2003), which is responsible for pseudouridylation, the first RNA modification found in tRNA and ribosomes (Cohn and Volkin., 1951; Charette and Gray., 2000). Pseudouridylation is thought to be required for modifying RNA structure by enhancing base-to-base stacking (Charette and Gray, 2000). TruB1 introduces pseudouridine at position 55 of tRNAs during the early stage of tRNA maturation (Becker et al., 1997; Zucchini et al., 2003). TruB1 mediated-pseudouridine is also found in mRNA (Schwartz et al., 2014; Carlile et al., 2014; Li et al., 2015; Safra et al., 2017). Notably, our CLIP analysis for endogenous TruB1 revealed a physical association of TruB1 with a series of mRNAs, although the functional significance of these interactions and the potential pseudouridylation were not determined. *E. coli* TruB has been reported to function as a chaperone for tRNA that is not mediated by its enzymatic activity (Keffer-Wilkes et al., 2016). This reflects our finding that let-7 maturation promoted by TruB1 is also independent of its intrinsic pseudouridylation enzyme activity. It would thus be interesting to test whether the interactions we observed between TruB1 and mRNA are also independent of its pseudouridylation enzyme activity.

Examination of the crystal structures of bacterial TruB and tRNA reveals that the base of U55 in the TUC stem-loop structure near position 55 of the tRNA is flipped to bind to the catalytic site of TruB (Hoang and Ferré-D’Amaré., 2001). In our HITS-CLIP, TruB1 was also bound to the stem-loop structure of let-7. Although the stem loop also exists in other miRNAs, only the let-7 stem loop is selectively regulated by binding of regulatory factors such as Lin28 and KSRP (Newman et al., 2008; Trabucchi et al., 2009). Consistent with this unique property of the let-7 the stem loop, TruB1 also recognizes this region, which may cause physical competition among these specific regulators. More detailed analyses are needed to determine the structural effects of TruB1 binding to let-7 and how it affects the microprocessing step of let-7 maturation.

As a cell biological function of the TruB1 and let-7 interaction, we found that TruB1 controlled cell proliferation by promoting let-7 maturation. The expression level of TruB1 correlates with cancer progression and/or oncogenesis in prostate cancer and pancreatic cancer, suggesting that TruB1 could be involved in the pathogenesis of cancer (Fig EV6B-D) (Edgar et al., 2002; Data ref: Edgar et al., 2002; Varambally et al., 2005; Data ref: Varambally et al., 2005; Pei et al., Data ref: Pei et al., Wang et al., 2015; Data ref: Wang et al., 2015). On the other hand, TruB2, which has the same enzyme activity site at TruB1, preferentially mediates pseudouridinylation of mitochondrial RNA, and is essential for cell survival by controlling glycolysis (Arroyo et al., 2016; Antonicka et al., 2017). TruB KO *E. coli* were weaker than wild-type cells when they were co-cultured; however this was not caused by decreased survival (Gutgsell et al., 2000). In this way, despite having highly conserved enzyme activity sites between species and homologs, their functions appear to be quite diverse.

PUS7, PUS10, DKC1, TruB1, TruB2, amongst others, are also pseudouridine synthases, and among them, PUS10 was recently reported to promote miRNA maturation also independently of its enzyme activity (Song et al., 2019). PUS10 introduces pseudouridine at position 55 of tRNA (Roovers et al., 2006), similar to TruB1, and position 54 of tRNA in archaea (Deogharia M, et al. RNA. 2019). PUS10 increases the affinity of microprocessors for various miRNAs and promotes their maturation (Song et al., 2019). However, these two pseudouridine synthases showed very different substrate selectivity. While TruB1 selectively and specifically interacts with pri-let-7 and promotes let-7 expression, PUS10 appears to non-specifically increase maturation of various miRNAs (Song et al., 2019). Consistent with this, our endogenous HITS-CLIP for TruB1, and the published PUS10 PAR-CLIP analysis, reveal that TruB1 binds to the terminal loop of let-7, while PUS10 binds to a site relatively far from the terminal loop of miRNAs (Song et al., 2019). Furthermore, TruB1 acts suppressively on cell proliferation due to the selective promotion of let-7 maturation, whereas PUS10 tended to promote cell proliferation (Song et al., 2019). Thus, TruB1 functions differently to PUS10.

TruB is known to be an RNA modifying enzyme that possesses two distinct molecular functions; RNA folding and subsequent RNA modification. There are other such examples of RBPs with dual functions in addition to their intrinsic enzymatic activity (i.e., ADRA1 / 2, METTL3, METTL16) (Ota et al., 2013; Lin et al., 2016; Pendleton et al., 2017). Our strategy and data reveal an unexpected function of TruB1 in miRNAs biogenesis with let-7 specificity, and present a new aspect of miRNA regulation by RNA binding proteins.

## Materials and Methods

### Plasmid construction

The pLuc2-KAP-MCS vector was generated by a previously reported method (Ito et al., 2017). To create the let-7 sensor vector, the chemically synthesized let-7 complementary sequence was annealed and inserted between the EcoRI and XhoI sites. We used the empty pLus2-KAP-MCS vector as the control sensor (−) vector. The KRAS mRNA vector was generated by inserting the let-7a target site of Kras mRNA into pLuc2-KAP-MCS at the EcoR1 and Kpn1 sites. The sequence encoding human TruB1 ORF was cloned by PCR from 293FT cell cDNA. pcDNA-TruB1 was generated by inserting the ORF of human TruB1 with a Flag sequence into pcDNA.3.1(+) (Thermo Fisher Scientific) at the Nhe1 and EcoR1 sites. The pCLT vector was designed and created by modifying pCS 2 (RIKEN) using the tet-on system. The sequence is shown in Appendix Table S5. pCLT-TruB1 and pCLT-GFP were generated by inserting the ORF of human TruB1 and GFP with a Flag sequence into the pCLT vector at the Nhe1 and EcoR1 sites. pCLT-mt1 and mt2 were generated by PrimeSTAR Mutagenesis Basal Kit (Takara). pET22b-TruB1, mt1 and mt2 were generated by inserting the ORF of human TruB1 and its mutants into the pET22b vector (Merck Millipore) at the NdeI and XhoI sites. pcDNA-pri-let-7a1 was generated by inserting the human pri-let-7a1 sequence into pcDNA.3.1(+) at the BamH1 and Not1 sites. Loop mt was generated by inverse PCR from pcDNA-pri-let-7a1. The nucleotide sequences of the primers used for the PCR are shown in Appendix Table S4.

### Cell culture

HEK293FT, A549 and HeLa cells were maintained in DMEM medium (Corning) supplemented with 10% FBS (Gibco) and 1% penicillin–streptomycin (Wako) at 37 °C with 5% CO_2_. siRNAs [(TruB1 (NM_139169.3), TruB2 (NM_015679.1), or negative controls (SN-1002); Bioneer), (LIN28B (s52477) or negative control (AM4611); Ambion, Life Technologies) and 2’-O-methylated antisense inhibitor (let-7 family or negative control: detailed sequences shown in Appendix Table S4, Nippon-shinyaku) were transfected into cells using Lipofectamine RNAiMAX (Thermo Fisher Scientific) using the Manufacturer’s recommended protocol. For transient overexpression, pcDNA vectors (pcDNA-TruB1-Flag) were transfected with FuGENE HD (Promega) into cells according to the Manufacturer’s instructions. For steady state expression, lentiviral vectors [pCLT vectors (pCLT-GFP, TruB1 and its mutants)] were used with doxycycline (Clontech) treatment (1.0 μg/ml) for 5 days. For virus production, HEK293FT cells were transfected with the lentiviral vector pCLTs together with the packaging plasmids pCMV-VSVG-RSV-Rev (addgene) and pCAG-HIVgp (addgene) at a 1:0.5:0.5 ratio using PEI Max reagent (Polysciences) according to the Manufacturer’s instructions. Supernatants were harvested 48 h after transfection. Cells (5 × 10^5^) were mixed in 2ml viral supernatant supplemented with 8μg/ml polybrene (Merck Millipore) and further incubated at 37°C. The infected cells were selected by 1 μg/ml puromycin (Thermo Fisher Scientific) and kept in selection medium for 7 days.

### Genome editing

TruB1-Flag tagged HEK293FT cells were established by using CRISPR–Cas9 system. ALT-R XT CRISPR RNA (crRNA) (Integrated DNA Technologies) and ALT-R trans-activating crRNA (tracrRNA) (Integrated DNA Technologies) were resuspended in Nuclease-free-duplex buffer (Integrated DNA Technologies) to a final concentration of 100 μM. crRNA and tracRNA with equal volumes were mixed and heated for 5 minutes at 95 °C, followed by gradually cooling down to room temperature. Mixed RNAs were incubated at room temperature for 20 minutes with ALT-R Cas9 Nuclease V3 (61 μM) (Integrated DNA Technologies) and single-stranded oligo DNA (ssODN) including sequences of HiBIT, 3 x Flag and complementary sequence of N-terminal in TruB1 genome. Next, this RNA, protein and ssODN were co-transfected into HEK-293FT cells using a NEPA21 Super Electroporator (NEPAGENE). We used the HiBIT system to detect the knocked-in cells (Schwinn et al., 2018) with dilution methods. After transfection, knocked-in cells were detected by Nano Glo HiBiT Lytic Detection System (Promega) using the Manufacturer’s recommended protocol. Knocked-in cells were verified by PCR and Western blotting with anti-Flag antibody (MBL).

### Cell-based screening

We created 384-well reporter library plates for cell-based screening. In detail, library plasmids were diluted to 4 ng/μL in elution buffer (10 mM Tris·HCl; pH 8.5) in 96-well plates and were then dispensed into 384-well culture plates at 5 μL per well. Cell-based screening for let-7 regulator genes was performed by modifying a previously described method (Ito et al., 2017). Five microliters of OPTI-MEM (Gibco) containing 0.15 μL of PEI-Max (Polyscience), 20 ng of sensor vector or control sensor (−) vector, and 5 ng of pRL-SV40 (Promega) Renilla luciferase construct was added to the 384-well reporter library plates and incubated for 20 min. Then 293FT cells (Thermo Fisher Scientific) in 40 μL of DMEM containing 10% of FBS were added into each well using an automated multidispenser ECO DROPPER III (AS ONE). The cells were cultured in a 5% CO2 incubator at 37 °C for 48 h. Luciferase activity was measured by means of an ARVO X3 (PerkinElmer) and the Dual-Glo Luciferase Assay System (Promega).

### Luciferase reporter assay

Luciferase assays were performed by incubating 20 μL of OPTI-MEM containing 0.4 μL of FuGENE HD, 25 ng of reporter plasmids and 5 ng of pRL-SV40 for 10 min in 96-well culture plates. Then 293FT cells in 100 μL of DMEM containing 10% of FBS were added to each well and cultured for 48 h. Luciferase activity was measured using an ARVO X3 (PerkinElmer) and the Dual-Glo Luciferase Assay System (Promega).

### Reverse transcription and RT-qPCR

Total RNA samples were extracted using ISOGEN (NIPPON GENE). To evaluate mRNA expression, reverse transcription was carried out using oligo dT primers (Thermo Fisher Scientific) and SuperScript III (Thermo Fisher Scientificic). Quantitative RT-PCR (qRT-PCR) was run using THUNDERBIRD SYBR qPCR Mix (TOYOBO). The results are expressed as mRNA levels normalized to GAPDH mRNA expression in each sample. To evaluate microRNA expression, reverse transcription was performed using the TaqMan MicroRNA Reverse Transcription Kit (Thermo Fisher Scientific) and TaqMan MicroRNA Assays (Thermo Fisher Scientific). Quantitative RT-PCR (qRT-PCR) was run using THUNDERBIRD Probe qPCR Mix (TOYOBO). The results are expressed as RNA levels normalized to RNU6B expression in each sample. The primer sequences and assays used for the real-time PCR are listed in Table S4. To determine RNA levels, at least three independent RNA samples were analyzed.

### Northern blotting

LNA/DNA probes (PAGE grade, fasmac, Table SX) were labelled with 32P-γ-ATP by means T4PNK (Takara), followed to filter by MicroSpin G-25 Columns (GE healthcare). 4 ug of small RNA purified by the mirVana miRNA Isolation kit (Thermo Fisher Scientific) was heated at 90 °C for 20 sec with gel loading buffer II (Thermo Fisher Scientific) and placed on ice, followed by running on a 15% TBE-Urea PAGE. RNA was transferred to a Hybond-N+ membrane (GE Healthcare) at 4mA/cm^2^ for 45 min and fixed by UV-crosslinking. The membrane was dried and pre-hybridized at 37 °C for 30 min in ExpressHyb (Clontech). Hybridization was performed in ExpressHyb containing RI-labeled probe at 37 °C overnight with rotation. Subsequently, the membrane was washed twice in low stringency buffer (2X SSC, 0.05% SDS) at room temperature for 30 min and twice in high stringency buffer (0.5X SSC, 0.1% SDS) at room temperature for 40 min. Membranes were exposed to an autoradiography film.

### TaqMan Array

Total RNA was isolated using the mirVana miRNA isolation kit (Thermo Fisher Scientific). 3μL of RNA (1000 ng) were reverse transcribed by Megaplex RT Primers, Pool A (human) (Thermo Fisher Scientific) using the TaqMan® microRNA Reverse Transcription Kit (Thermo Fisher Scientific). For qRT-PCR, RT product was mixed with TaqMan® Universal PCR Master Mix, No AmpErase UNG (Thermo Fisher Scientific) and the appropriate amount of water, and subsequently loaded into the ports of the TaqMan array cards. RealTime PCR was performed on a 7900HT Fast Real-Time PCR System with TaqMan® Array Block (Thermo Fisher Scientific), using universal cycling conditions (95 °C/10 min, then [95 °C/15 sec, 60 °C/60 sec] for 40 cycles). Raw Ct values were normalized to RNU6B.

### Western blotting

Total protein samples were extracted by radioimmunoprecipitation assay (RIPA) buffer [50 mM Tris·HCl (pH 8.0), 150 mM NaCl, 0.5% deoxycholate (DOC), 0.1% SDS, 1% Nonidet P-40]. Proteins in the cell lysates were separated by SDS/PAGE followed by semidry transfer to a PVDF membrane. Membranes were blocked for 60 min with Blocking One (Nacalai Tesque), incubated with an anti-Flag (M185-3L, MBL), anti-DGCR-8 (ab90579, abcam), anti-Lin28B (11965S, Cell signaling technologies) or anti– β-actin (A5316; Sigma Aldrich) at 4 °C overnight, rinsed, and then incubated for 1 h with ECL mouse IgG HRP-conjugated whole antibody (GE Healthcare) or rabbit IgG HRP-conjugated whole antibody (GE Healthcare). The blot was developed with the ECL Select Western Blotting Detection Reagent (GE Healthcare). The protein levels are normalized to β-actin levels in each sample.

### HITS-CLIP

Flag-tagged cells were collected and crosslinked with 254 nm UV-crosslinking at 400 mJ/cm2 and 200mJ/cm2 on ice in a Ultraviolet Crosslinker (UVP). The cells were collected, centrifuged and resuspended in Lysis buffer (1xPBS, 0.1% SDS, 0.5% DOC, 0.5% NP-40). After incubation with rotation for 10 min at 4°C, cells were sonicated 30sec/30sec (on/off) for 5cycles by Bioruptor (Cosmo BIO). Then, whole cell lysate was collected by centrifugation at 13,000 × g for 15 min at 4 °C and treated with 5 U/ml RNase T1 (Thermo Fisher Scientific) and Turbo DNase (Thermo Fisher Scientific) for 5 min at 37°C. Three different doses of RNaseT1 and one non-crosslinked cells were used as controls. For immunoprecipitation, we used mouse anti-FLAG antibody (MBL) and normal mouse IgG (Sigma Aldrich) as an additional negative control. Antibodies were coupled to Dynabeads Protein G (Thermo Fisher Scientificc) following the Manufacturer’s protocol. Coupled antibodies were washed with PBST. RNaseT1 treated whole cell lysate and coupled antibodies were mixed and rotated at 4°C for 2h. Antibody-bound proteins and RNA-protein complexes were isolated by magnetic stand and washed with wash buffer (1xPBS, 0.1% SDS, 0.5% DOC, 0.5% NP-40) and high-salt-wash buffer (5xPBS, 0.1% SDS, 0.5% DOC, 0.5% NP-40). Purified RNA fragments were treated with CIP (Takara) and ligated with 3’ linker adaptor by using T4 RNA ligase (Thermo Fisher Scientific) at 16°C overnight. Treated RNA fragments were labelled with 32p-γ-ATP, followed by electrophoresis with SDS-PAGE (NuPAGE, Thermo Fisher Scientific) and transferred to nitrocellulose membranes. RI-labelled RNAs were detected by autoradiography. Appropriate bands or smear-bands were picked up from the membrane and treated with Proteinase K (Roche). RNA fragments were purified by Acid phenol :CHCL3 (Thermo Fisher Scientific). Purified TruB1-bound RNA fragments were ligated with 5′ linker RNA, followed by the reverse-transcriptase reaction using RT primers and SuperScriptIII (Thermo Fisher Scientific). cDNA libraries were amplified by means of Phusion High-Fidelity DNA Polymerase (New England Biolabs). Quantification of libraries was calculated using Quibit3.0 (Thermo Fisher Scientific), TapeStation (Agilent) with High Sensitivity DNA Kit, and qPCR analysis by NEB Next Library Quant Kit for Illumina (New England Biolabs). All sequencing was performed using an Illumina MiSeq (Illumina) with MiSeq Reagent Kit v3 (Illumina).

### HITS-CLIP analysis

Raw reads (fastq file) obtained from the Illumina pipeline were checked, trimmed and clipped using Bioconductor, FastQC, FASTX-toolkit and cutadapt (v.1.2.1) and were subsequently mapped to the human genome (build hg38) using bowtie2. To eliminate reads in which the adaptor was ligated more than once, adaptor removal was performed 3 times. Duplicate reads were filtered by IGV tools. All data analysis was done using a combination of custom and publicly available tools (featureCounts, SAMtools, RSeQC, bedtools, tRNAscan-SE 2.0 and R) including resources available from the UCSC Genome Browser and Galaxy Bioinformatics (http://main.g2.bx.psu.edu/). All aligned reads shorter than 10nt that mapped to more than a single position in the genome were excluded from further analysis. In trimmed data, the 500-bp flanking sequences of known pre-miRNAs were extracted as pri-miRNAs. Transcript annotation was acquired from UCSC. The read count is plotted for miRNAs with TaqMan array data in this report. Data was normalized by expression levels of each microRNA in control samples of the TaqMan array.

### Motif Analysis

We identified motifs for sequences mapped to tRNAs. Reads were randomly selected within 5000 reads by seqkit (https://bioinf.shenwei.me/seqkit/) to perform the following analysis. Multiple Em for Motif Elicitation (MEME) was performed using v.4.9.1 of the MEME browser application.

### RNA immunoprecipitation (RIP)

Cells were collected, centrifuged and resuspended in Lysis buffer (1×PBS, 0.1% SDS, 0.5% DOC, 0.5% NP-40). After incubation with rotation for 10 min at 4°C, cells were sonicated 30sec/30sec (on/off) for 5 cycles. Then, whole cell lysate was collected by centrifugation at 13,000 × g for 15 min at 4 °C. For immunoprecipitation, we used mouse anti-FLAG (MBL), anti-DGCR-8 (abcam) and normal mouse IgG (Sigma Aldrich) as an additional negative control. Antibodies were coupled to Protein G Dynabeads (Thermo Fisher Scientific) following the Manufacturer’s protocol. Coupled antibodies were washed with PBST. Whole cell lysate and coupled antibodies were mixed and rotated at 4°C for 2h. Antibody-bound proteins and RNA-protein complex were isolated by magnetic stand and washed with wash buffer (1xPBS, 0.1% SDS, 0.5% DOC, 0.5% NP-40) and high-salt-wash buffer (5xPBS, 0.1% SDS, 0.5% DOC, 0.5% NP-40). RNA was eluted by Acid phenol :CHCL3 (Thermo Fisher Scientific). Pri-miRNA levels were analyzed by qRT-PCR. Relative ratio of protein bound pri-miRNA was calculated as compared to its input sample.

### In vitro processing assay

pri-miRNAs were in vitro transcribed using MEGA script T7 Transcription Kit (Thermo Fisher Scientific) with a 32p-α-UTP (RI) label, followed by PAGE purification. We also synthesized RI-labeled pri-let-7a1 in which pseudouridine was randomly introduced using UTP: pseudouridine (Carbosynth) at a ratio of 1:1. Cells were collected, centrifuged and resuspended in 200 μL of Lysis buffer (WCL) (20mM Tris-HCL(pH 8.0), 100mM KCL, 0.5mM EDTA, 5% glycerol, 0.5mM PMSF and 5mM DTT). After incubation with rotation for 10 min at 4 °C, cells were sonicated 15sec/90sec (on/off) for 3 cycles. Whole cell lysate was collected by centrifugation at 18,000 × g for 15 min at 4 °C. Labelled RNA was treated with whole cell lysate and 6.4mM MgCL_2_ at 37 °C for 90 min. RNA was eluted by Acid phenol :CHCL3 (Thermo Fisher Scientific), and run on 15% TBE-Urea PAGE. RNA was visualized by autoradiography after gel drying.

### Electrophoretic mobility shift assay (EMSA)

pri-miRNAs were in vitro transcribed using MEGA script T7 Transcription Kit (Thermo Fisher Scientific) with 32p-α-UTP to label by RI, followed by PAGE purification. Labelled RNA and recombinant proteins were mixed and incubated at 37 °C for 1 h, followed by naïve PAGE. RNA and RNA-protein complexes were visualized by autoradiography.

### Expression and purification of recombinant proteins

*E. coli* BL21 (DE3) (Novagen) was transformed by the plasmids, and the transformants were grown at 37 °C until the A600 reached 1.0. The expression of the TruB1 proteins was induced by adding isopropyl-β-D thiogalactopyranoside at a final concentration of 0.1 mM and incubating the cultures for 18 h at 18 °C. The cells were collected and lysed by sonication in buffer, containing 20 mM Tris-HCl, pH 7.0, 500 mM NaCl, 10 mM β-mercaptoethanol, 20 mM imidazole, 0.1 mM phenylmethylsulfonyl fluoride, and 5% (v/v) glycerol. The proteins were first purified by chromatography on a Ni-NTA agarose column (QIAGEN), and then further purified on a HiTrap Heparin column (GE Healthcare). Finally, the proteins were purified by chromatography on a HiLoad 16/60 Superdex 200 column (GE Healthcare), in buffer containing 20 mM Tris-HCl, pH 7.0, 200 mM NaCl, and 10 mM β-mercaptoethanol. The purified proteins were concentrated and stored at − 80 °C until use. The hTruB1 mutants were purified in a similar manner.

### In vitro enzymatic activity assay

pri-miRNAs were in vitro transcribed using MEGA script T7 Transcription Kit (Thermo Fisher Scientific) with 32p-α-UTP, followed by PAGE purification. Labelled RNA was treated with recombinant TruB1 or its mutants in reaction buffer (20mM TrisHCL (pH 8.0), 0.1mM EDTA, 100mM NaCl, 5mM DTT and 100mM NH4Cl) at 37 °C for 60 min. RNA was eluted by Acid phenol :CHCL3 (Thermo Fisher Scientific), and treated with Nuclease P1 (New England BioLab) at 37 °C for 3 h. RNA was run on 1D-thin layer chromatography on cellulose plates (TLC, 20 cm, Merck Millipore) with solvent (isopropanol/HCl/water (70/15/15); v/v/v) for 6 h at room temperature. RNA was visualized by autoradiography.

### CMC Primer extension assay

Primers (PAGE grade, fasmac, Appendix Table S4) were labelled with 32P-γ-ATP by T4PNK (Takara), followed by filtering through MicroSpin G-25 Columns (GE healthcare). 4 ug of total RNA was treated with 0.17M N-cyclohexyl-N0 -(2-morpholinoethyl)carbodiimide metho-p-toluenesulfonate (CMC, Tokyokasei) in reaction buffer (7M urea, 50mN Bicine and 4mM EDTA) at 37 °C for 20 min. Reaction was stopped with 100 μL of stop buffer [0.3 M NaOAc and 0.1 mM EDTA, pH 5.6] and 700 μL of EtOH. After incubation at −80 °C for 1 h. The treated RNA was centrifuged, and washed twice with 70% EtOH. RNA pellet was dissolved in 50mN Na2CO3 (pH 10.4) and incubated at 37 °C for 4 h. Reaction was stopped in the same manner. RNA pellet was dissolved in water and purified using the RNeasy MinElute Cleanup kit (QIAGEN). Treated RNA was reverse transcribed with RI-labelled primer by SuperScript III (Thermo Fisher Scientific) with or without ddATP solution (Thermo Fisher Scientific). cDNA was eluted by phenol/chloroform/isoamyl alcohol (Thermo Fisher Scientific), and run on 10% TBE-Urea PAGE. cDNA was visualized by autoradiography after gel drying.

### Evaluation of cell viability

Cell viability was determined using RealTime-Glo MT Cell Viability Assay (Promega). A total of 1000 cells/well were plated in 96-well culture plates. After 24 h of incubation, the medium was changed to a growth medium with MT Cell Viability Substrate (Promega) and Nano Luc Enzyme (Promega). Then, the cells were incubated for an additional 0, 12, 24, 48, 72 h of culture. The luminescence in the resulting solution was measured using an ARVO X3 (PerkinElmer) at each point. Cell viability was determined by dividing the luminescence activity by that of the control cells. These experiments were conducted on at least three independent samples.

### Data analysis from public databases

We browsed several series (GEO accession GSE3325, GDS4102, GSE64333_at; Varambally et al., 2005; Pei et al., Wang et al., 2015) of the GEO Profiles database (Edgar et al., 2002).

## QUANTIFICATION AND STATISTICAL ANALYSIS

Blot signals were quantified using ImageJ (NIH) (Schneider et al., 2012). Data are presented as the means ± standard errors, and differences between groups were evaluated using the Student’s t-test. A P-value of <0.05 was considered statistically significant. Asterisk in figures indicate differences with statistical significance as follows: *p < 0.05. All the experiments were conducted in close adherence to the institutional regulations.

## Data Availability

All the data required to reproduce this study are included in this published article and supplementary information. The raw data of TaqMan array and HITS-CLIP were deposited in the GEO under accession number GSE143508 and GSE143510, respectively. Further information and requests for resources and reagents should be directed to and will be fulfilled by Hiroshi Asahara.

## Acknowledgements

We thank Shuji Takada, Satoshi Hara and Asuteka Nagao for technical assistance and support, and Yutaro Uchida, Hiroki Tsutsumi, Maiko Inotsume, Haruka Hosogai, Hiroto Yamamoto and all other members of the Department of Systems BioMedicine in Tokyo Medical and Dental University for their support, and Helen Pickersgill at Life Science Editors for scientific critical reading and editorial assistance. This research was supported by AMED-CREST from AMED (Japan Agency for Medical Research and Development) (JP19gm0810008 to H.A.), JSPS KAKENHI (Grant numbers: 17K15018, 19K16794 to R.K. 18H03980 to K.T. 26113008, and 15H02560, 18K19603, 19KK0227 to H.A.), and grants from the National Institutes of Health (Grant numbers: AR050631, AR065379) to H.A.

## Author contributions

R.K., T.C., Y.I., T.M., K.M., T.S., K.T., and H.A. designed the study; R.K. performed experiments, analyzed data; K.T. created and purified recombinant proteins; R.K., T.C., and Y.Y. generated sequencing data. T.S. and K.T. supervised the project; R.K., T.S., K.T., and H.A. drafted the manuscript; H.A. critically reviewed the manuscript; and all authors approved the final manuscript.

## Expanded View Figure Legends

**Figure EV1. Schematic model of cell-based functional screen**

Repressors of maturation such as Lin28 lead to high luciferase activity, and activators lead to low luciferase activity. This cell-based screen is assaying for a function directly on endogenous let-7 maturation, rather than affinity/binding. The gain-of-function screen also enables to identify regulatory factors that may cause lethality under loss of function conditions, such as with Crispr-Cas9 or RNAi-KD screens.

**Figure EV2. Knockdown of TruB1 repressed expression of relative miRNAs in various human cells.**

(A). qRT-PCR analysis for TruB1 mRNA expression normalized to GAPDH. TruB1 KD by siRNA significantly repressed the expression of TruB1 mRNA in HEK-293FT cells. Error bars show SD; n=3. (B, C). qRT-PCR analysis for miRNAs expression in A549 cells (B) and Hela cells (C) with TruB1 KD. TruB1 KD by siRNA significantly repressed the expression of the let-7 family. Error bars show SD; n=3.

**Figure EV3. Impact of pseudouridylation enzyme activity on microprocessing of let-7.**

(A). EMSA of 32p-ATP-labelled tRNA ^Phe^ mixed with recombinant TruB1, mt1 or mt2 at several doses. RNP: Ribonucleoprotein complexes. (B). Protein expression of Flag-tagged TruB1 and its mutants. Protein expression was evaluated by Western blotting. Protein was isolated from HEK-293FT cells infected with tetracycline-inducible lentiviruses expressing TruB1, mt1, or mt2, 5 days after doxycycline treatment. (C). Location of pseudouridine sites were detected by CMC primer extension. Total RNA purified from HEK-293FT cells was treated with CMC. CMC-treated RNA were reverse-transcribed with RI-labelled specific primers for tRNAphe or pri-let-7a1. ddATP was used as a sequence control. (D,E). In vitro processing analysis for pri-let-7 with pseudouridine. 32p-ATP-labelled pri-let-7a1 was synthesized using UTP : pseudouridine at a ratio of 1:1 or 1:0. This labelled RNA was treated with whole cell lysate from TruB1 expressing HEK293FT cells transfected with pcDNA3.1-TruB1. Autoradiographed image (D) and the relative processing rate (E) are shown.

**Figure EV4. loop mutant of pri-let7a1 and HITS-CLIP for TruB1.**

(A). Loop sequence of mutant RNA in which the loop structure of pri-let-7a1 was modified (loop mt).

(B). Protein expression of 3 x Flag-tagged TruB1 in genome edited cells. Flag-tagged TruB1 was most highly expressed in ED-9 cells.

(C). Western blotting for input and IP samples in HITS-CLIP.

(D). Autoradiography of RI-labelled RNA in a complex with TruB1 (black frame).

(E). Agilent data of cDNA library show smeared band at 150 – 250 bp.

(F). Mapped reads of sequenced data to human genome.

**Figure EV5. Relative pri-let-7a1 expression in HEK-293 cells with knockdown of Lin28B.**

qRT-PCR analysis for pri-let-7a1 expression normalized to GAPDH. Lin28B KD by siRNA tended to increase the expression of pri-let-7a1, but this was not statistically significant. Error bar shows SD; n=3.

**Figure EV6. TruB1 suppressed tumor progression.**

(A). TruB1 and mt1 suppressed cell proliferation. Real-time glo assay for HEK-293 cells infected with tetracycline-inducible lentiviruses expressing TruB1, mt1, or GFP, 5 days after doxycycline treatment. TruB1 and mt1 suppressed cell proliferation. Error bars show SD; n=3.

(B-D). Relationship between TruB1, let-7, and tumor progression in various cancers.

(B). Published microarray and microRNA-sequencing in prostate biopsy samples from patients with prostate cancer (data accessible at NCBI GEO database (Edgar et al., 2002), accession GSE64333, Wang et al., 2015; Data ref: Wang et al., 2015). In these data, there were correlations between the expression levels of TruB1 and both let-7b and let-7i. Regarding let-7a, a statistically significant correlation was not found, but the same tendency was observed. (C, D). TruB1 inversely correlates with tumor progression or oncogenesis in prostate cancer (C) and in pancreatic cancer (D). LD: local disease. Data from (Varambally et al., 2005; Data ref: Varambally et al., 2005; Pei et al., Data ref: Pei et al., 2009)

## Appendix table title and legends

## Appendix tableS1. Raw data of cell-based screening

## Appendix tableS2. Raw data of TaqMan array

## Appendix tableS3. Mapped miRNA and tRNA in HITS-CLIP

(A) Tag densities were representative as table. Density of miRNA clusters in HITS-CLIP were normalized by miRNA expression as determined by TaqMan array. (B). Mapped tRNA in HITS-CLIP.

## Appendix tableS4. Sequences of oligonucleotide, DNA and RNA used in this paper

## Appendix tableS5. Sequences of pCLT vector.

